# Towards a comprehensive delineation of white matter tract-related deformation

**DOI:** 10.1101/2021.04.13.439136

**Authors:** Zhou Zhou, Xiaogai Li, Yuzhe Liu, Madelen Fahlstedt, Marios Georgiadis, Xianghao Zhan, Samuel J. Raymond, Gerald Grant, Svein Kleiven, David Camarillo, Michael Zeineh

**Affiliations:** Department of Bioengineering, Stanford University, Stanford, CA, 94305, USA; Neuronic Engineering, KTH Royal Institute of Technology, Stockholm, 14152, Sweden; Department of Radiology, Stanford University, Stanford, CA, 94305, USA; Department of Neurosurgery, Stanford University, Stanford, CA, 94305, USA; Department of Neurology, Stanford University, Stanford, CA, 94305, USA; Department of Mechanical Engineering, Stanford University, Stanford, CA, 94305, USA

**Keywords:** Axonal injury, Tract-related deformation, Injury criteria, Finite element analysis, Computational brain modelling

## Abstract

Finite element (FE) models of the human head are valuable instruments to explore the mechanobiological pathway from external loading, localized brain response, and resultant injury risks. The injury predictability of these models depends on the use of effective criteria as injury predictors. The FE-derived normal deformation along white matter (WM) fiber tracts (i.e., tract-oriented strain) has recently been suggested as an appropriate predictor for axonal injury. However, the tract-oriented strain only represents a partial depiction of the WM fiber tract deformation. A comprehensive delineation of tract-related deformation may improve the injury predictability of the FE head model by delivering new tract-related criteria as injury predictors. Thus, the present study performed a theoretical strain analysis to comprehensively characterize the WM fiber tract deformation by relating the strain tensor of the WM element to its embedded fiber tract. Three new tract-related strains with exact analytical solutions were proposed, measuring the normal deformation perpendicular to the fiber tracts (i.e., tract-perpendicular strain), and shear deformation along and perpendicular to the fiber tracts (i.e., axial-shear strain and lateral-shear strain, respectively). The injury predictability of these three newly-proposed strain peaks along with the previously-used tract-oriented strain peak and maximum principal strain (MPS) were evaluated by simulating 151 impacts with known outcome (concussion or non-concussion). The results preliminarily showed that four tract-related strain peaks exhibited superior performance than MPS in discriminating concussion and non-concussion cases. This study presents a comprehensive quantification of WM tract-related deformation and advocates the use of orientation-dependent strains as criteria for injury prediction, which may ultimately contribute to an advanced mechanobiological understanding and enhanced computational predictability of brain injury.

## Introduction

Traumatic brain injury (TBI) is a growing public health concern worldwide. In the United States, the number of TBI-related victims in 2013 is estimated to be 2.8 million, resulting in 2.5 million emergency department visits, 282000 hospitalizations, and 56000 deaths.^1^ A meta-analysis of injury data from 16 European countries reported an overall TBI incidence rate of 262 per 100000 and an average fatality rate of 11 per 100000.^2^ Olesen and colleagues^3^ estimated a total cost of 33 billion Euros associated with TBI in 2010 in Europe. Despite global endeavors to prevent the occurrence and alleviate the consequence of TBI, neither a clear decrease in TBI-induced mortality nor improvement of overall outcome has been observed.^4^ This is particularly true for axonal injury, which is notoriously underreported,^5^ difficult to diagnose,^6^ and associated with immediate and persistent impairment to attention and cognition.^7^ This TBI-related urgency indicates that the current understanding of TBI pathogenesis is insufficient and cannot provide solid guidance for the development of head protection strategies and therapeutic tactics.

As numerical surrogates, finite element (FE) models of the human head have been increasingly used to explore the mechanobiological pathway from external loading, localized brain response, and resultant injury risk. The injury predictability of such numerical surrogates requires accurate anatomical and mechanical representations of each intracranial structure, precise descriptions of the interaction among various intracranial components, and appropriate criteria as injury predictors.^8^ In part by using advanced neuroimaging techniques, it is feasible for current FE models to incorporate a high degree of structural detail. For example, several three-dimensional (3D) models possess a sophisticated anatomical depiction of the lateral ventricles^9, 10^ and the skull,^11, 12^ by leveraging magnetic resonance imaging (MRI) and computed tomography scans, respectively. In parallel, persistent endeavors in experimental characterization of biological tissue guide the establishment of constitutive equations for numerical implementation. Recently, Mihai and coworkers^13^ presented a family of hyperelastic models with material constants calibrated against the experimental curves of brain tissue under finite simple shear superposed on varying axial tension or compression.^14^ Emerging improvement has also been noted in the modeling of intracranial interfaces, especially for the fluid-structure interaction (FSI) scenarios. Recently, arbitrary Lagrange-Eulerian and smoothed particle hydrodynamics approaches were respectively employed to emulate the brain-skull interface and brain-ventricle interface, contributing to an enhanced representation of the intracranial fluid dynamics.^15–18^ However, contemporary head models appear to lack comparable explorations of the different aspects of brain injury criteria.

While promising progress has been noted in narrowing down the knowledge gaps between the external loading and brain responses globally or regionally with the aid of finite element head models,^19–23^ great disparities exist among models when interpreting the estimated responses by means of injury criteria. Various measures characterizing the local mechanical responses of brain tissue have been derived from the computational studies to assess brain injury risk, such as intracranial pressure,^24^ stress,^25, 26^ strain,^19–21, 27^ etc. Even when limited to strain-based metrics, the adopted criteria vary from maximum principal strain (MPS),^28^ maximum shear strain,^10^ to strain-based derivatives (e.g., strain rate,^29^ the product of strain and strain rate,^30^ and cumulative strain damage measure (CSDM)^31^). No consensus has been reached yet on the best metric to describe tissue deformation and discriminate injury severity. Considering that axonal injury is a type of trauma in which diffuse lesions occur within fiber tracts usually consisting of oriented axons, any measures that account for the orientation of fiber tracts are anatomically relevant and may function as promising candidate criteria. White matter (WM) fiber orientation and distribution from larger bundles can be derived from an MRI technique called diffusion tensor imaging (DTI).^32^ In DTI, the fiber orientation in each voxel is represented by a rank-2 symmetric tensor, visually depicted by an ellipsoid.^33^ Recent studies have integrated the orientation information of fiber tract delineated by DTI into the FE model to extract the deformation pertaining to fiber tracts.^9, 30, 34–55^ Across these numerical endeavors, a specific component, i.e., tract-oriented strain measuring the normal deformation along the WM fiber tracts, was extracted by resolving the strain tensor along the fiber orientation. Of note, the tract-oriented strain is synonymous with axonal strain or fiber strain.^43, 48, 50^ It has been repeatedly reported by several independent groups that the maximum tract-oriented strain exhibited improved injury predictability than MPS.^30, 43, 45, 46, 52, 55^

Despite the inspiring initial results, the tract-oriented strain only represents a partial depiction of WM fiber tract deformation. From a mathematical perspective, projecting the deformation (described by a strain tensor) along the fiber orientation to attain the tract-oriented strain (described by a scalar) discards a large amount of information originally conveyed by the strain tensor. A comprehensive delineation of tract-related deformation that appropriately utilizes the full information of the strain tensor may hold the potential to further improve injury predictability of FE models. Such potential is supported by the findings of *in vitro* models.^56–59^ For example, through spatiotemporal correlation between neurite pathomorphology and the deformation of neurons, Bar-Kochba and colleagues^56^ found that bleb formation was strongly correlated with shearing along the longitudinal direction of neurons, while the timeframe of neuronal death was related to the integral of the stretching over the length of the neuron. Nakadate and coworkers^59^ delivered uniaxial stretching to a two-dimensional (2D) culture slice with randomly oriented axons and noted swellings in axons with their orientations exhibiting various angles relative to the stretching direction. Based on the above analyses, any criteria related to the deformation of WM axonal fibers may be candidate predictors for axonal injury, the full array of which have yet to be properly considered in FE head models.

The aim of this study is to present a comprehensive delineation of WM fiber tract deformation and investigate the injury predictability of novel tract-related strain metrics. To achieve this, a theoretical strain analysis was performed by relating the strain tensor of the WM element to its embedded fiber tracts. Exact analytical solutions of three new tract-related strain measures were presented, characterizing the normal deformation perpendicular to the fiber tracts, and shear deformation along and perpendicular to the fiber tracts, respectively. The injury predictability of these three newly-proposed strain peaks along with the previously-used maximum tract-oriented strain and MPS was evaluated by simulating 151 impacts with known outcome (concussion or non-concussion). We hypothesize that the four tract-related strain peaks exhibit superior injury predictability than MPS.

## Methods

### Finite element head model

The FE model (Fig. 1) used in this study was previously developed at the KTH Royal Institute of Technology in Stockholm^28^ using the LS-DYNA software. Responses of the model have shown good correlations with experiments of brain-skull relative motion,^60, 61^ intracranial pressure,^22^ and brain strain.^62^ The model includes the scalp, skull, brain, subarachnoid cerebrospinal fluid (CSF), ventricles, dura mater, falx, tentorium, pia mater, eleven pairs of the largest parasagittal bridging veins, and a simplified neck with the extension of the spinal cord. Anatomical differentiation was implemented to divide brain elements into cerebral gray matter (GM), cerebral WM, corpus callosum, thalamus, brainstem, midbrain, and cerebellum (Fig. 1B-C). Detailed information regarding the geometry discretization and material choice for each head component, as well as interface conditions among various intracranial structures, are available in previous studies.^15, 28^ Particularly, because of the lack of consensus regarding the mechanical anisotropy and heterogeneity of the brain^63–65^ the brain was simulated as a homogeneous and isotropic medium with its nonlinear behavior described by a second-order hyperelastic constitutive law with additional linear viscoelastic terms to account for its rate dependence.^28^

**Fig. 1.**
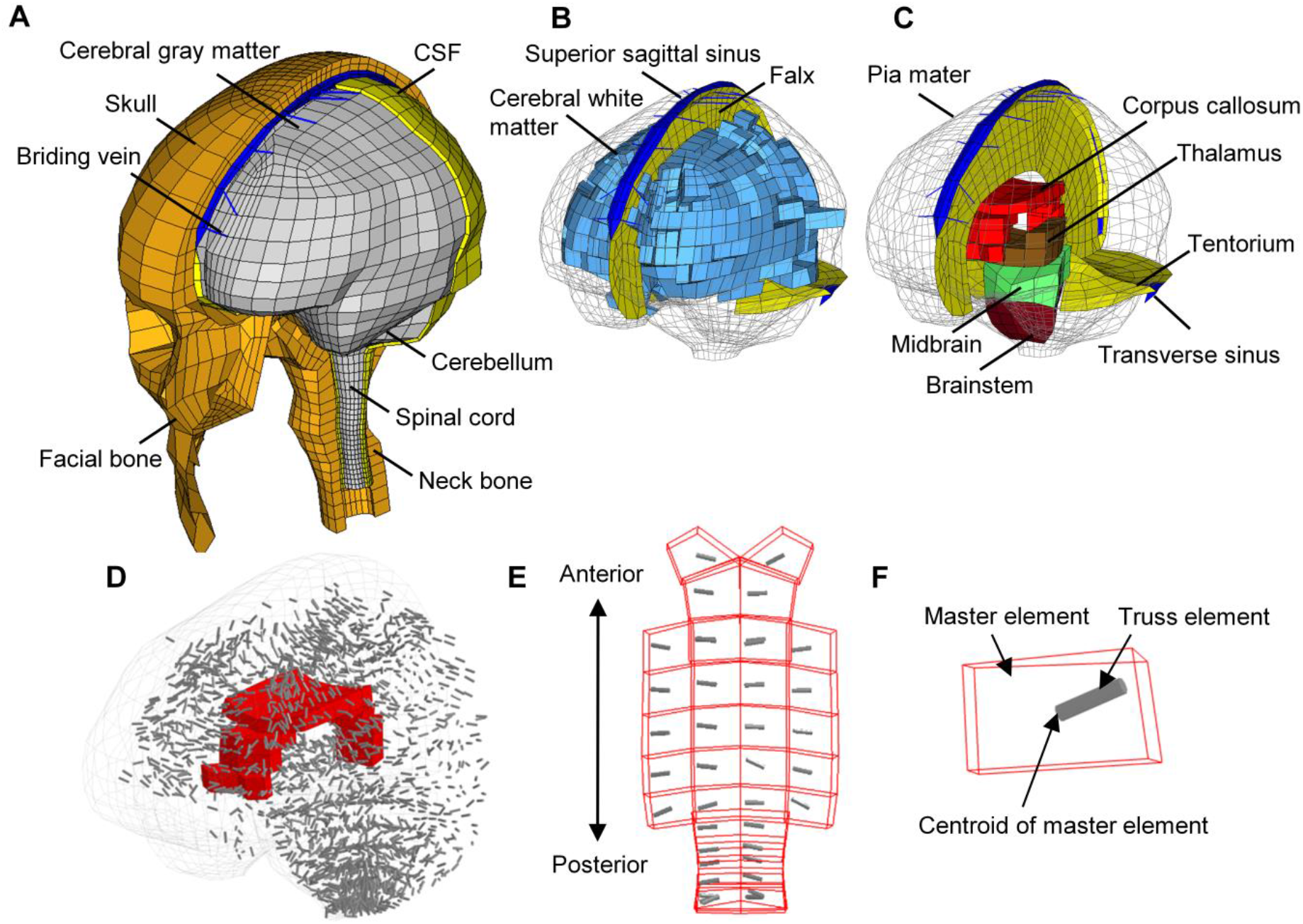
Finite element model of the human head with embedded truss elements. (**A**) Finite element model of the human head with the skull open to expose the CSF and brain. (**B**) Model of cerebral white matter, superior sagittal sinus, and falx. (**C**) Model of deep brain structures, pia mater, tentorium, and transverse sinus. (**D**) Brain with embedded truss elements representing the dynamically-changing local fiber orientation, and the corpus callosum elements in red. (**F**) Top view of the truss elements within the corpus callosum. (**E**) A representative element in the corpus callosum with a truss element embedded.

### Embedded element method for real-time fiber orientation

Given that the deformation of WM fiber tracts is directly dependent on the real-time fiber orientation, an embedded element method was implemented in the FE model to monitor the temporal fiber orientation (Fig. 1D-F) following the approach presented earlier.^42, 54^ Briefly, the brain mesh of the head model was voxelized to generate a reference volume, which was aligned to the volume of the ICBM DTI-81 atlas^32^ via an affine registration. To insert the orientation information delineated by DTI with a resolution of 1 mm to the FE model with a mean element size of 5.2 mm, a weighted averaging procedure was applied to partially compensate for the resolution mismatch. For each brain element in the head model, corresponding voxels in the DTI atlas that fell inside the given element were identified. Diffusion parameters (i.e., diffusion tensor and FA value) of these identified voxels were averaged using equations (1) and (2), through which the diffusion information from those voxels closer to the centroid of the element was more heavily weighted:

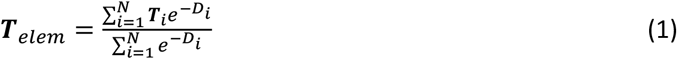

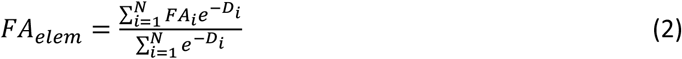

where ***T**_elem_* and *FA_elem_* are respectively the mean diffusion tensor and mean FA value calculated for each brain element; *N* is the number of selected voxels for a given brain element; ***T**_i_* and *FA_i_* are respectively the diffusion tensor and FA value of each selected voxel; *D_i_* is the distance from the voxel to the element centroid.

In the current study, those brain elements with the *FA_elem_* values over 0.2 were considered as WM with its fiber orientation approximated as the first eigenvector of ***T**_elem_*. This orientation information was concretely represented by truss elements (serving as slave elements), which were embedded within the WM elements (serving as master elements). Note that the embedded truss elements only served as auxiliaries for obtaining the updated orientation of fiber tracts at each time step without additional contribution to mechanical stiffness. For a given truss element, one node was located at the centroid of its master element, while the other fell within the boundary of its master element (Fig. 1F). Both acceleration and velocity of a given truss element were determined exclusively by its master element without relative motion between them. Consequently, the real-time fiber orientation during head impacts was reflected by the temporal direction of the truss element and was updated at each timestep of the simulation. Details regarding the coupling protocol between DTI and WM element and the implementation of the embedded element method for tracking the real-time fiber orientation are previously reported by Giordano and colleagues^42^ and Zhou and coworkers,^54^ respectively.

### Derivation of tract-related strains via coordinate transformations

#### Computation of tract-related strains at a given timestep

To decipher WM fiber tract-related deformation at a specific timestep, a theoretical strain analysis was performed via coordinate transformations,^66^ in which the strain tensor of WM element was transformed to the coordinate systems with one axis aligned with the real-time fiber orientation. As illustrated in Fig. 2, both the Green-Lagrange strain tensor for a representative WM element and the orientation of its embedded truss element for a given timestep were extracted from a pre-computed simulation and then transformed into the skull-fixed coordinate system (***XYZ***) (Fig. 2B) with ***X*** = [1 0 0]^*T*^, ***Y*** =[0 1 0]^*T*^, and ***Z*** = [0 0 1]^*T*^. The strain tensor and the fiber orientation within the skull-fixed coordinate system are denoted as ***E***_skull_ and ***ν***. The ***ν*** is a unit vector with its three components along the ***X***, ***Y***, and ***Z*** axes as *ν*_1_, *ν*_2_, and *ν*_3_ respectively. Details about the transformation of the strain tensor from the global coordinate system to the skull-fixed coordinate system are available in the previous publications from our group^47, 54^ as well as others.^30^

**Fig. 2.**
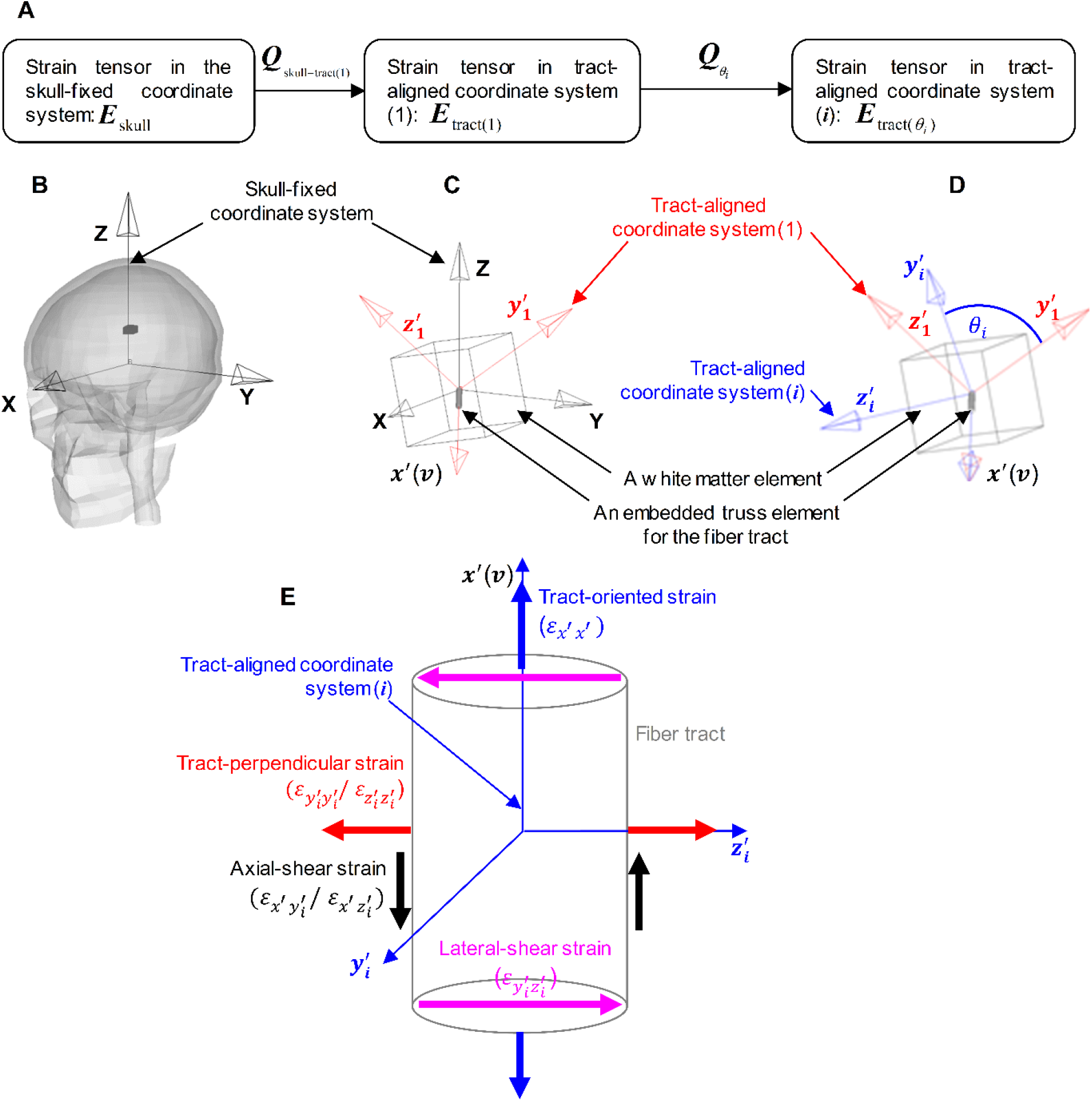
Demonstration of coordinate systems used in the theoretical analysis via the coordinate transformations to delineate the tract-related deformation for one representative WM element at one timestep. (**A**) Summary of the coordinate transformations of the strain tensor illustrated in subfigure (**B**)-(**D**). (**B**) An FE model of the human head with a representative white matter element highlighted in dark gray. A skull-fixed coordinate system and corresponding axes are illustrated with the origin at the head’s center of gravity. (**C**) A representative white matter element with an embedded truss element representing the tract orientation. A modified skull-fixed coordinate system with the origin at the centroid of the white matter element and the tract-aligned coordinate system (1) is superimposed. (**D**) The tract-aligned coordinate system (*i*) with the **x**′ axis aligned with the real-time tract orientation (***ν***). (**E**) Definition of tract-related strains with respect to the fiber tract in the tract-aligned coordinate system (*i*).

Given that the deformation of the fiber tracts is dependent on the fiber orientation, a specific tract-aligned coordinate system, i.e., the tract-aligned coordinate system (*1*) (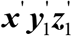) in Fig. 2C, was established with the origin at the centroid of the WM element and the ***x***′ aligned with the ***ν***. The 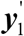 was computed as the cross product of ***x***′ and ***X***, while the 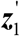 as the cross product of ***x***′ and 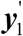.

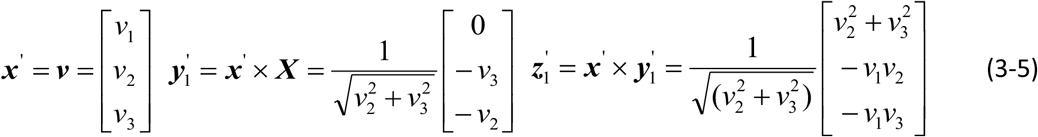

The strain tensor within the tract-aligned coordinate system (*1*), denoted by ***E***_tract(1)_, can be calculated as

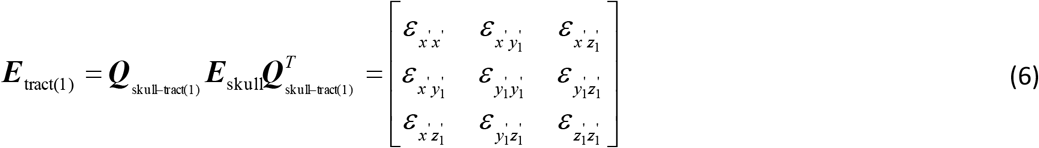

with the ***Q***_skull-tract(1)_ given by

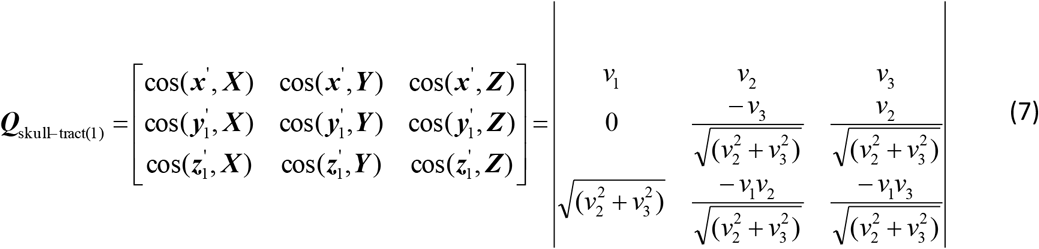

Since ***E***_skull_, *ν*_1_, *ν*_2_, and *ν*_3_ were output from LS-DYNA solver, all components of ***E***_tract(1)_ were known variables.

Within the context of having the ***x***′ aligned with the ***ν***, the ***x***′ was unique and the orthonormal unit vectors 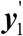 and 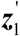 were instead non-unique. For all the components in the strain tensor, only *ε_x′x′_*, was exclusively dependent on the ***x***′, and, thus, can be obtained as

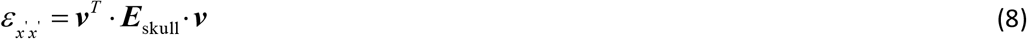

As depicted in Fig. 2E, *ε_x′x′_* is referred to as tract-oriented strain, characterizing the normal deformation along fiber tracts. The tract-oriented strain has been similarly extracted in previous studies.^9–30, 34–55^

For the remaining components in the transformed strain tensor, whose magnitudes are affected by the choice of 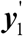 and 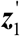, a new tract-aligned coordinate system, i.e., the tract-aligned coordinate system (*i*) 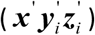 was introduced (Fig. 2D). In the tract-aligned coordinate system (*i*), the ***x***′ remained aligned with the ***v***, while the orthonormal unit vectors 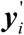 and 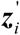 were arbitrarily established within the plane defined by 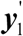 and 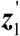. Note that, in the current study, the index *i* is commonly added as a subscript to identify all the variables related to the tract-aligned coordinate system (*i*) (e.g,. 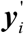, 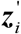), except for ***x***′ and *ε_x′x′_* that were constant across all tract-aligned coordinate systems. To quantify the angle between 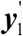 and 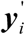 (equivalent to the angle between the 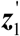 and 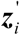), a new variable, *θ_i_*, was introduced (Fig. 2D). Thus, the strain tensor within the tract-aligned coordinate system (*i*) (***E***_tract(*θ*_*i*_)_) can be calculated as

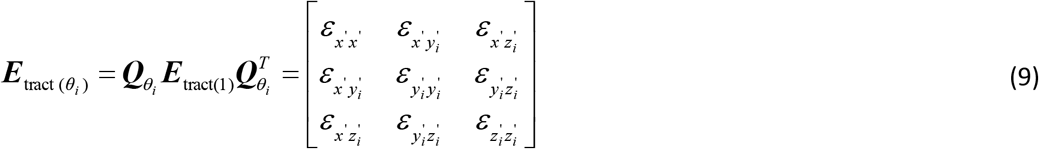

with the ***Q***_*θ*_*i*__ given by

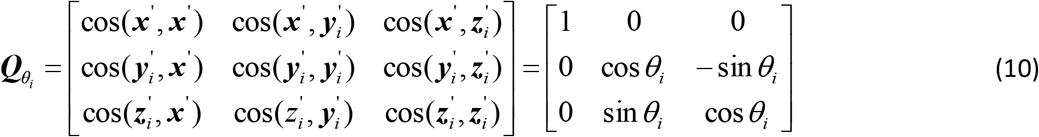

where 0° ≤ *θ_i_* ≤ 360°.

Integrating the equation (6) and equation (10) into equation (9), all components in ***E***_tract(*θ*_*i*_)_) can be obtained as a function of *θ_i_*.

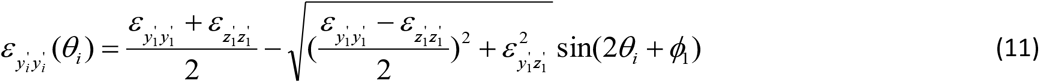

where 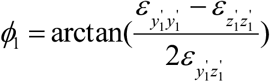.

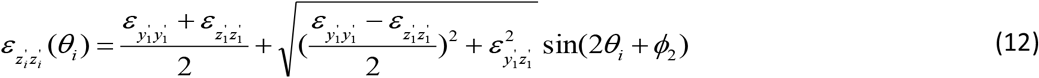

where 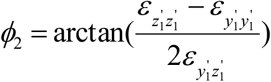.

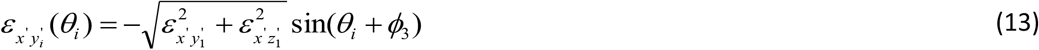

where 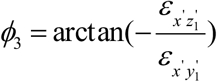.

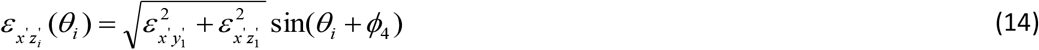

where 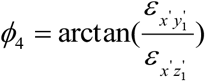.

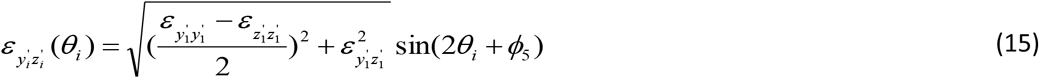

where 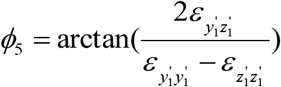.

Note that *ε_x′x′_* is exclusively dependent on the choice of ***x***′. Thus, *ε_x′x′_* remains constant across all tract-aligned coordinate systems and was expressed by equation (8).

In the current study, the peak values of equations 11-15 across the range of *θ_i_* were considered. The analytical solutions for the peak values for the 5 strain components as well as peak conditions are summarized in Table 1. Since the peak values of 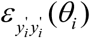 and 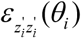 are identical, these two variables were combined as one and further referred to as tract-perpendicular strain (Fig. 2E),^57^ measuring the normal deformation perpendicular to fiber tracts. Similarly, the peak values of 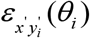 and 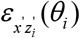 were combined as one variable and further referred to as axial shear strain (Fig. 2E),^56^ quantifying shear deformation along the fiber tracts. The peak value of 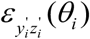 was termed lateral-shear strain (Fig. 2E),^57^ describing the shear deformation perpendicular to the fiber tracts. Consequently, once the strain tensor and real-time fiber orientation are known for a given timestep, the four tract-related strains can be calculated directly using the analytically derived formulas (Table 1).

**Table 1.** Analytical formulas for the tract-related strains described by equation 8 and equation 11-15 under a given time instance, and the respective conditions where the values were attained.

| Strain | Symbol | Peak value | Peak condition |
| --- | --- | --- | --- |
| Tract-oriented strain | $\varepsilon_{x'x'}$ | $\mathbf{v}^T \cdot \mathbf{E}_{\text{skull}} \cdot \mathbf{v}$ | Constant across all tract-aligned coordinate systems |
| Tract-perpendicular strain | $\varepsilon_{y_i y_i}(\theta_i)$ | $\frac{\varepsilon_{y_1 y_1} + \varepsilon_{z_1 z_1}}{2} + \sqrt{\left(\frac{\varepsilon_{y_1 y_1} - \varepsilon_{z_1 z_1}}{2}\right)^2 + \varepsilon_{y_1 z_1}^2}$ | $\sin(2\theta_i + \phi_1) = -1$ |
| | $\varepsilon_{z_i z_i}(\theta_i)$ | $\frac{\varepsilon_{y_1 y_1} + \varepsilon_{z_1 z_1}}{2} + \sqrt{\left(\frac{\varepsilon_{y_1 y_1} - \varepsilon_{z_1 z_1}}{2}\right)^2 + \varepsilon_{y_1 z_1}^2}$ | $\sin(2\theta_i + \phi_2) = 1$ |
| Axial-shear strain | $\varepsilon_{x' y_i}(\theta_i)$ | $\sqrt{\varepsilon_{x' y_1}^2 + \varepsilon_{x' z_1}^2}$ | $\sin(\theta_i + \phi_3) = -1$ |
| | $\varepsilon_{x' z_i}(\theta_i)$ | $\sqrt{\varepsilon_{x' y_1}^2 + \varepsilon_{x' z_1}^2}$ | $\sin(\theta_i + \phi_4) = 1$ |
| Lateral-shear strain | $\varepsilon_{y_i z_i}(\theta_i)$ | $\sqrt{\left(\frac{\varepsilon_{y_1 y_1} - \varepsilon_{z_1 z_1}}{2}\right)^2 + \varepsilon_{y_1 z_1}^2}$ | $\sin(2\theta_i + \phi_5) = 1$ |

#### Computation of tract-related strains across the entire simulation

The derivation described above was extended to all timesteps across the entire simulation. This derivation assumed that the displacements across the fiber tracts and surrounding matrix were continuous. The yielded analytical solutions are consistent with previous results in the *in vitro* models for special cases of simple shear^57, 58^ and compression.^56^

### Loading condition

Simulations of clinically diagnosed head impacts were conducted, through which strain-based measures were obtained. At Stanford University, instrumented mouthguards have been developed to measure six-degree-of-freedom head kinematics during in-game head impacts in athletes.^67, 68^ Using these instrumented mouthguards, over 500 head impacts in football have been tracked and video-confirmed.^69^ In the current study, two concussed impacts (i.e., one with the athlete suffering alteration of consciousness, and the other with the player had a milder but self-reported concussion) and 113 randomly selected non-concussed impacts were simulated.^69^ All simulated loadings were collected from collegiate American football players at practices and training events. Each impact trace possessed a recorded duration of 100 ms.

To further enrich the loading cases, 53 football impacts in a recently analyzed database of National Football League (NFL) collisions^70^ were simulated. The impact kinematics of these NFL collisions sustained by the professional football players were obtained from laboratory reconstruction using Hybrid III anthropometric test dummies.^71, 72^ While the loading traces of these 53 simulated cases exhibited varying durations ranging from 34 ms to 100 ms, a case-by-case scrutinization was performed following the requirement that the maximum strains be attained within the available impact durations. Only 36 impacts, including 15 concussive impacts and 21 non-concussive impacts, met the requirement and were retained for further analysis.

In summary, a total of 151 impacts (including 17 concussed impacts and 134 non-concussed impacts) were considered from the two datasets (Fig. 3). In each simulation, the skull was assumed to be rigid, and the translational and rotational accelerations were prescribed to a node located at the center of gravity of the FE head model and rigidly attached to the skull. A selectively reduced integration scheme was used for all the brain components and CSF. A typical simulation of 90 ms required about 11 hours using LS-DYNA 11.0 with double precision on a local Linux cluster with a single processor. The model responses were output at every 0.5 ms.

**Fig. 3.**
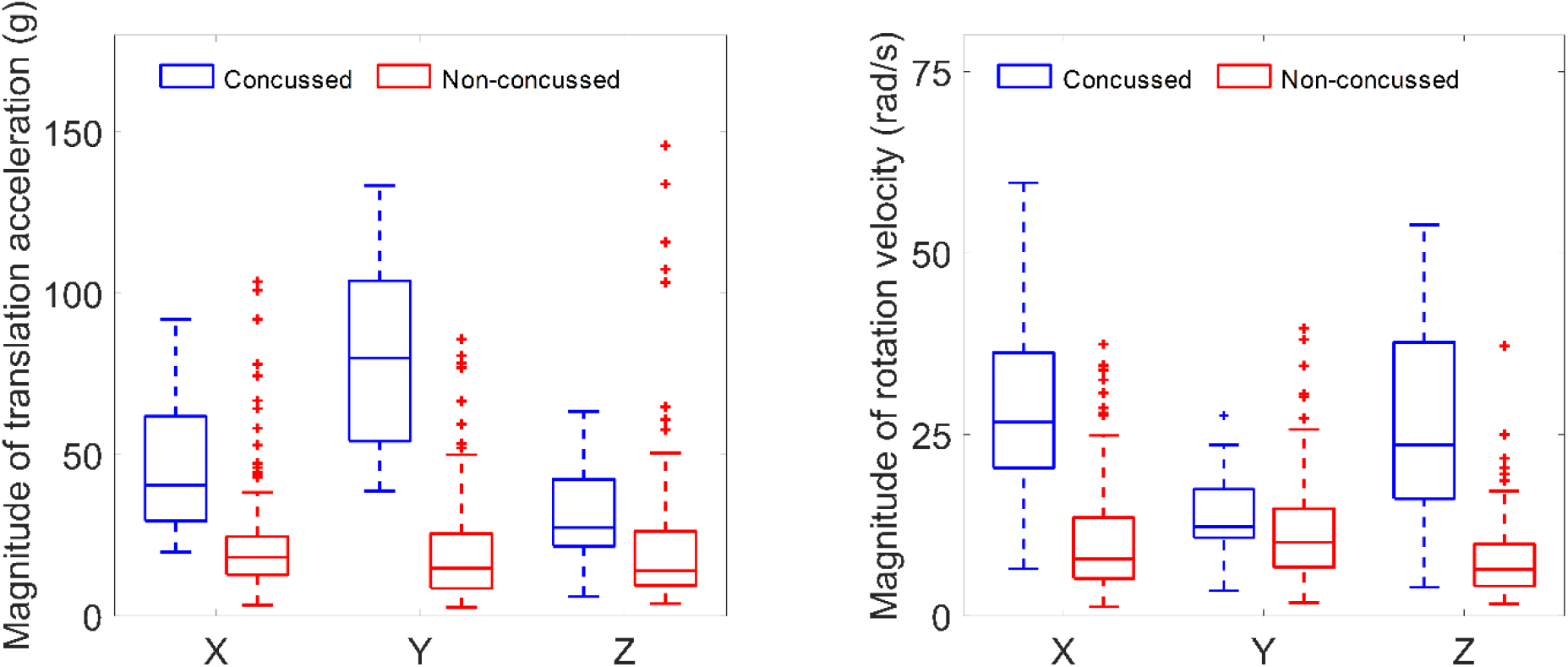
Boxplot of kinematic magnitudes of the concussed (N=17, blue) and non-concussed (N=134, red) impacts. Note the X, Y, and Z axes are the same as those in the skull-fixed coordinate system in Fig. 2B. On each box, the central line is the median value, and the upper and lower edges of the box are the 25^th^ and 75^th^ percentile values, while the outliers are shown as ‘+’ symbol outside the box.

### Strain-based injury metrics

The peaks of the four tract-related strains across the entire simulations were proposed as injury metrics (Table 2), i.e., maximum tract-oriented strain (MTOS), maximum tract-perpendicular strain (MTPS), maximum axial-shear strain (MASS), and maximum lateral-shear strain (MLSS). This is based on the rationale that the peak strain represents the worst scenario and is believed to cause injury to axonal fiber tracts. The injury predictability of the 4 tract-related strain peaks along with the prevalently-used MPS (Table 2) was evaluated. For these five strain-based metrics, the 95^th^ percentile maximum values in five commonly injured sub-regions (i.e., cerebral WM, brainstem, midbrain, corpus callosum, and thalamus) and the whole WM were obtained. In light of the common practice of extracting MPS at the entire brain level,^73–76^ the predictability of MPS extract from the whole brain elements was additionally examined. Thus, the injury predictability of these 5 strain-based metrics was evaluated either at the regional level or whole WM/brain level, similar to previous studies.^28, 43, 52, 77^

**Table 2.** Summary of the abbreviation of the metrics evaluated in this study and the corresponding description.

| Abbreviation | Description |
| --- | --- |
| MPS | The peak of the 1<sup>st</sup> principal Green-Lagrange strain across the entire simulation. |
| MTOS | The peak of normal strain along the white matter fiber tracts across the entire simulation. |
| MTVS | The peak of normal strain perpendicular to the white matter fiber tracts across the entire simulation. |
| MASS | The peak of shear strain along the white matter fiber tracts across the entire simulation. |
| MLSS | The peak of shear strain perpendicular to the white matter fiber tracts across the entire simulation. |

### Statistical analysis

While the four tract-related strains are fundamentally different measures according to their analytical formulas in Table 1, the interdependence between the MPS and four tract-related strain metrics remained unclear. To ascertain this, we conducted linear regression analyses between MPS and each of four tract-related strains, respectively. The tests were performed based on the element-wise peak values across the entire simulation for all WM elements (N=1323) from the simulation of a representative concussive impact (i.e., Case 157H2 from the NFL dataset). In addition to the correlation coefficient (r), the root mean square error normalized by the range of MPS (i.e., NMSRE) was also calculated for each analysis.

We next evaluated the injury predictability of 5 strain-based metrics (Table 2) in terms of discriminating the concussive and non-concussive cases. To correlate the binary injury outcome (i.e., concussion or non-concussion) with the continuous injury metrics (95^th^ percentile values), survival analysis with a Weibull distribution was employed (N=151). The survival analysis treats all data as double censored, producing a model that is more appealing on physical groups, while a Weibull distribution always satisfies a zero risk of injury for zero force. This statistical model was similarly adopted in previous biomechanical studies^46, 55^ with the corresponding merits previously detailed by McMurry and Poplin.^78^ To further assure an objective comparison and maximize the training dataset, all survival analyses were conducted within a leave-one-out cross-validation (LOOCV) framework.^79, 80^ To evaluate the sensitivity and specificity of each strain-based metric to separate the concussion and non-concussive cases, the receiver operating characteristic (ROC) curve analysis was generated both for the testing and training datasets based on a threshold of 50% injury probability. The area under each ROC curve (AUC) was calculated with a higher value indicating better injury predictability and a value of 0.5 for a random guess.

All the statistical analyses were performed with custom scripts in MATLAB (R2016b, MathWorks, Natick, MA) with built-in statistical toolbox functions and SPSS (22.0, SPSS Inc. Chicago, Illinois). The threshold for significance was p<0.05.

## Results

### Case illustration

To demonstrate the applicability of the proposed analytical formulas in Table 1 to computing the tract-related strains and the extraction of injury metrics in Table 2, we employed Case 157H2 from the NFL dataset as a representative impact for illustration purposes. Case 157H2 is a concussive impact in which the football player was laterally struck, resulting in high angular velocities in the coronal and sagittal planes. The impact duration was 50 ms. Note that, in the following two sections, the computation tract-related strain for Case 157H2 is firstly illustrated by one representative element in the corpus callosum and then extended to all WM elements.

#### Computation of tract-related strains for one element

As exemplified by one randomly selected element in the corpus callosum in Fig. 3, the first principal strain was directly output from the LS-DYNA solver for each timestep. To acquire the tract-related strains, both the strain tensor of the exemplified element and the temporal orientation of the embedded truss element were extracted from the pre-computed simulation and then transformed within the skull-fixed coordinate system. Hence, four strain-based measures pertaining to the fiber tracts were computed using the analytical formulas in Table 1 at each timestep. The peaks of these four tract-related strain measures along with the first principal strain over the entire simulation were considered as potential injury metrics in the current study. For this specific element, the value (occurring time) of each metric is highlighted in Fig. 4.

**Fig. 4.**
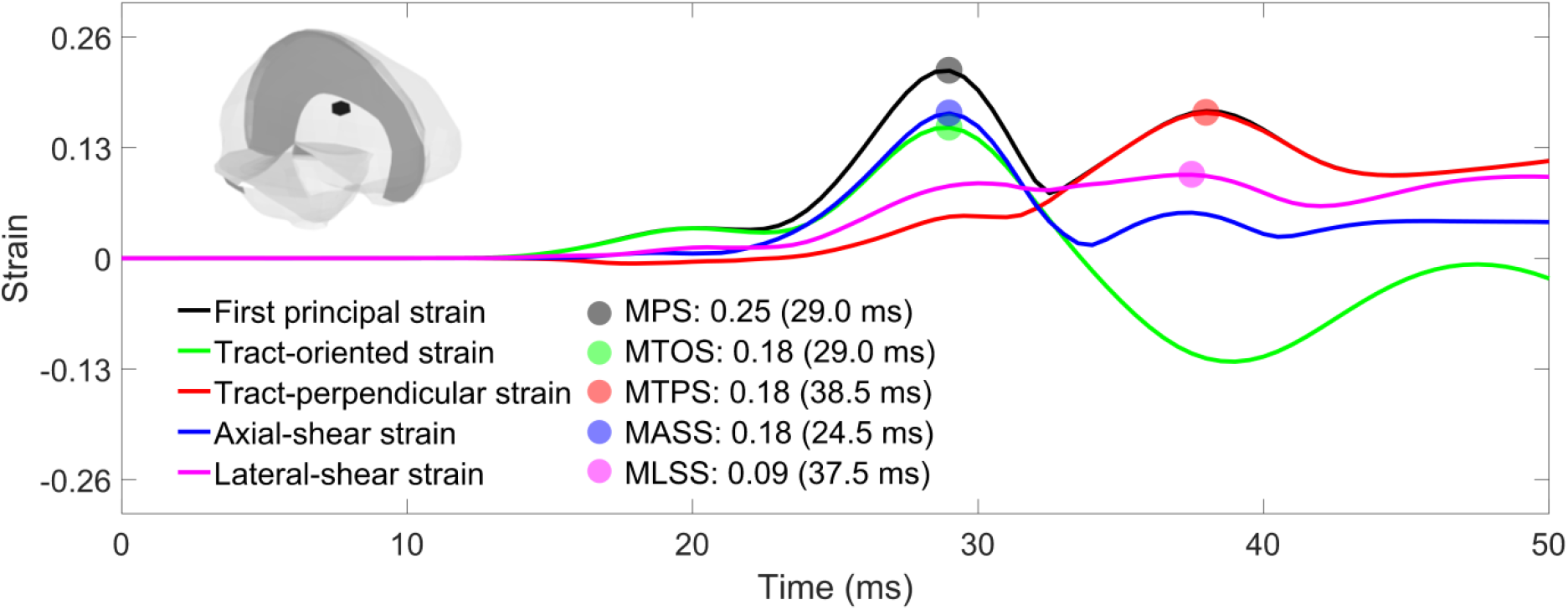
Time history curves of strain-based measures for one representative element in the corpus callosum (highlighted in black within the whole-brain thumbnail in the top-left corner) during a simulation of a concussed impact (Case 157H2). The peak values for all curves were highlighted as filled dots that were considered as injury metrics in the current study.

#### Computation of tract-related strains for one impact

The calculation detailed above for one representative element was applied to all WM elements (N=1323) in the FE model for the illustrated case. The median values of element-wise strain peaks were 0.16 for MPS, 0.09 for MTOS, 0.11 for MTPS, 0.11 for MASS, and 0.09 for MLSS (Fig. 5A). For each of the 5 strain metrics, those WM elements with their strain peaks ranked within the top 25^th^ percentile were identified. As shown in Fig. 5B, disparities in spatial distribution and extent of WM regions predicted to endure high strains existed among the five strains. To further evaluate the interdependence between MPS and its tract-related counterparts, linear regression analyses were conducted between MPS and each of the four tract-related metrics, based on the results of all WM elements. Although a p-value less than 0.001 was attained for all four tests, the correlation coefficients (r) between the MPS and tract-related strains were between 0.43 and 0.73, with a NRMSE of 10.07-13.43% (Fig. 5C), indicating the tract-related strains cannot be trivially attained via scaling the MPS.

**Fig. 5.**
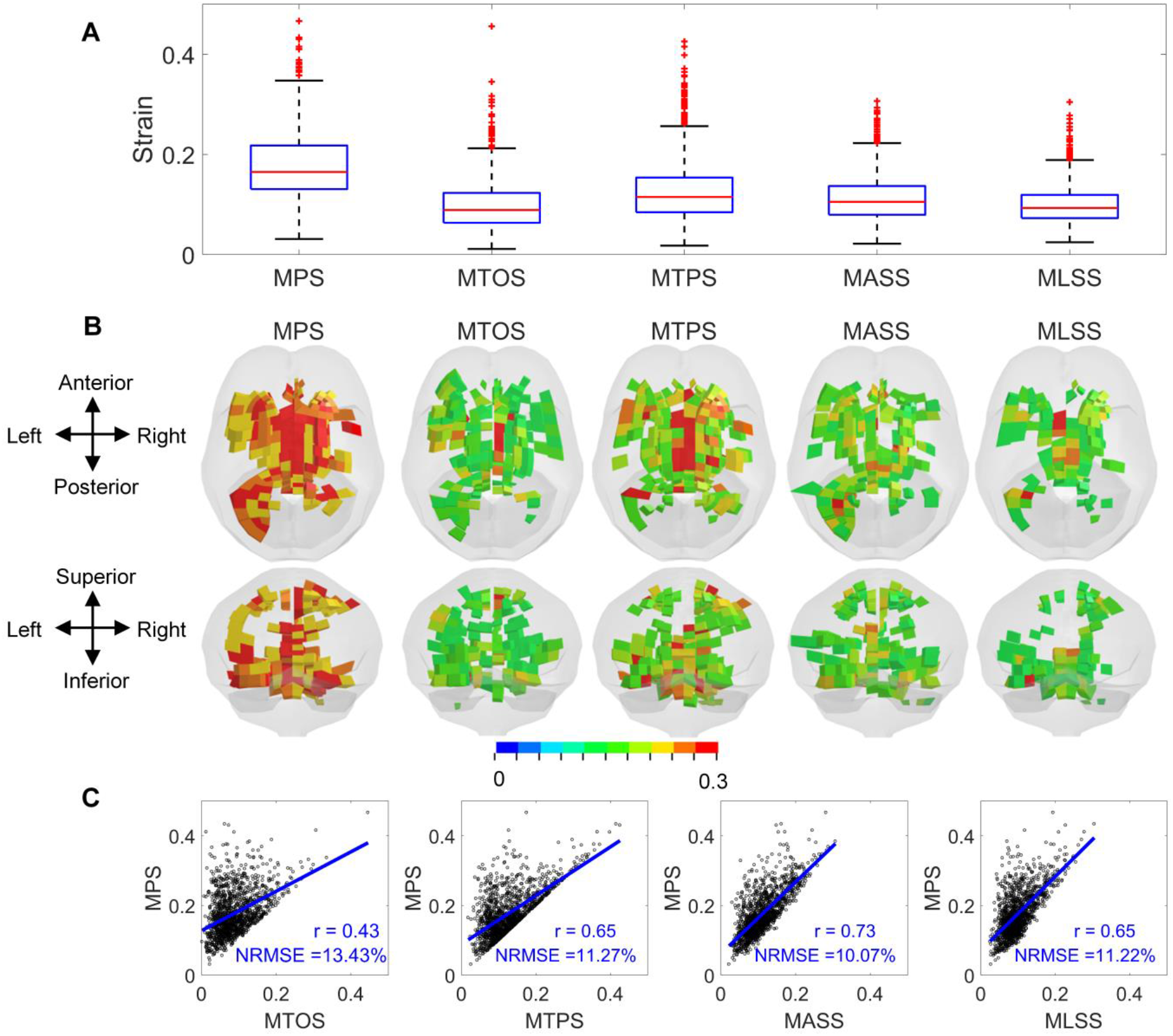
Model-estimated results based on 5 strain-based metrics in a representative impact involving one striking player with a concussion (Case 157H2) showing independence of the metrics. (**A**) Box plot showing element-wise strain-based metrics for all white matter elements (N = 1323). On each box, the central line (red) is the median value, and the upper and lower edges (blue) of the box are the 25th and 75th percentile values, while the outliers are shown as ‘+’ symbol (red) outside the box. (**B**) For each of the 5 strain-based metrics, white matter elements with strain peaks over the 75 percentile values calculated at the whole white matter level are identified and shown in axial and coronal views, respectively. For better illustration, the pia mater (gray) is additionally shown in translucency. (**C**) Linear regression plot for element-wise strain peaks between MPS and each of the 4 tract-related strains, respectively. A p-value of less than 0.001 was attained for all tests.

### Injury predictability of strain-based metrics

We next evaluated the discrimination performance of 5 strain-based metrics in terms of separating the concussive and non-concussive cases. A summary of the model-estimated metrics for different ROIs is presented in Fig. 6. The 5 strain-based metrics observed in the simulations are illustrated in the form of boxplot for concussed and non-concussed players in the cerebral WM, brainstem, midbrain, corpus callosum, thalamus, and whole WM (Fig. 6A-E) with the MPS additionally extracted from the whole brain shown in Fig. 6A. A statistically significant difference (Mann-Whitney U test, p<0.001) between concussed and non-concussed players was attained for all 5 metrics across all 5 subregions and the whole WM level (Fig. 6A-E). Similarly, the MPS value at the whole-brain level extracted from the concussed impacts was significantly different from that of the non-concussive impacts (Fig. 6A).

**Fig. 6.**
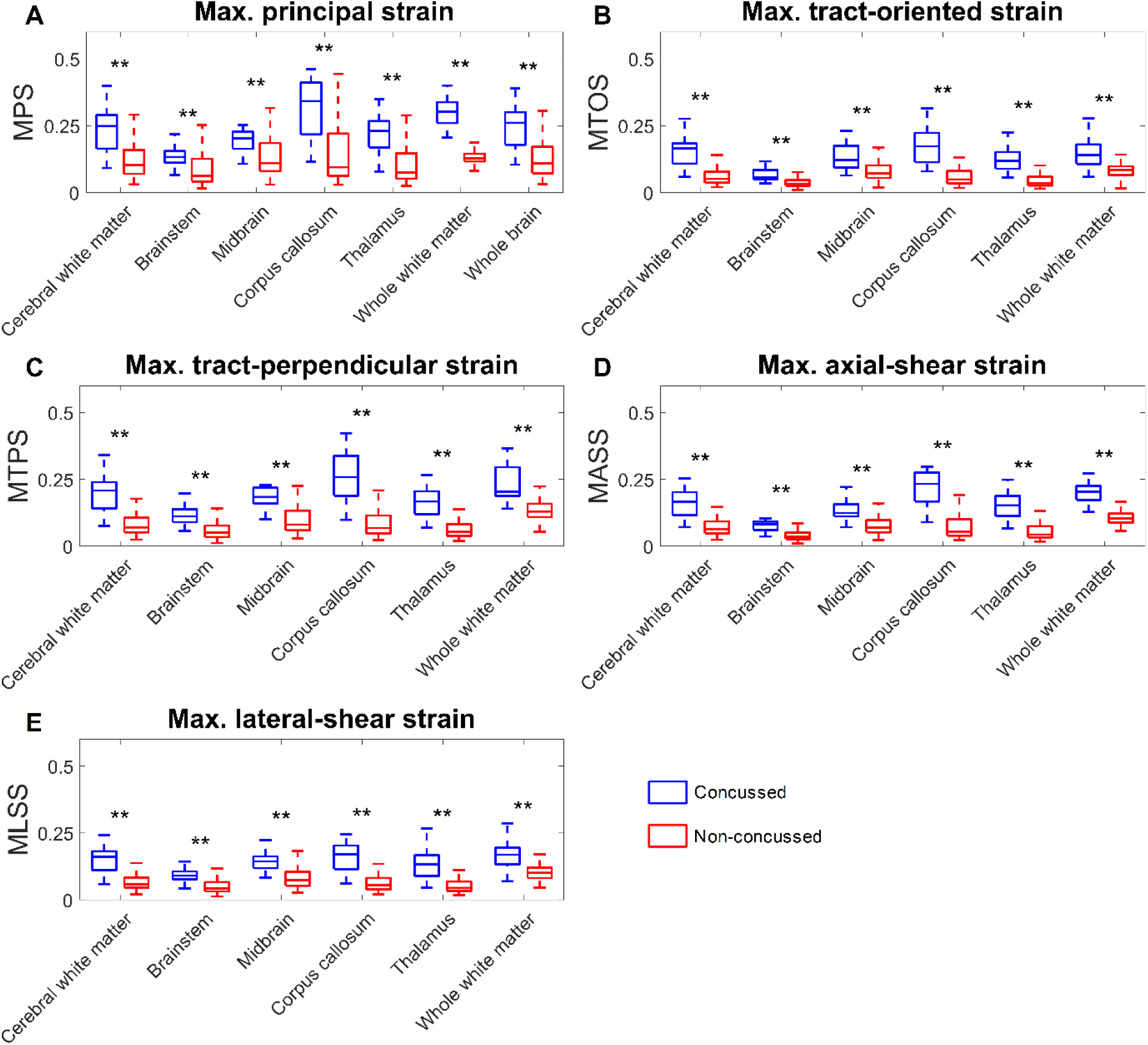
Regional 95^th^ percentile strain-related metrics. The values are shown in blue for concussed (n=17) and red for non-concussed (n=134) players for various anatomical regions in the brain model. On each box, the central line is the median value, and the upper and lower edges of the box are the 25^th^ and 75^th^ percentile values, while the error bars depict the total range of the data with the outlier being discarded. Statistical differences in Mann-Whitney U test are reported (**p<0.001). Note that subfigure (**A**) has an extra box depicting MPS results at the whole-brain level, in addition to the 5 regions being compared in subfigures (**B**)-(**E**).

The injury predictability of the five strain-based metrics in discriminating concussed from non-concussed cases was evaluated via survival analysis with the performance quantified by AUC values within the LOOCV framework. Fig. 7 shows the ROC curves for different strain-based metrics in different ROIs for the testing dataset. The AUC values for all metrics in all ROIs were over 0.5 (the AUC value for a random guess). As shown Fig. 7F, the AUC values for the four tract-related metrics at the whole WM level were 0.843 for MTOS, 0.869 for MTPS, 0.849 for MASS, 0.861 for MLSS, all of which were larger than those of MPS evaluated at the whole WM level (AUC = 0.825) and whole-brain level (AUC = 0.834). Similar trends were also noted in the five subregions (Fig. 7A-E) that the AUC values of all four tract-related metrics were consistently larger than that of MPS across all ROIs, while the metrics with the highest AUC values were region-dependent. For the cerebral WM, the MLSS exhibited the highest AUC value as 0.874, while MASS for brainstem (AUC = 0.868), MTPS for midbrain (AUC = 0.867), MTOS for corpus callosum (AUC = 0.909) and thalamus (AUC = 0.901). Similar findings were revealed by the training dataset with detailed results in Fig. A1 in Appendix.

**Fig. 7.**
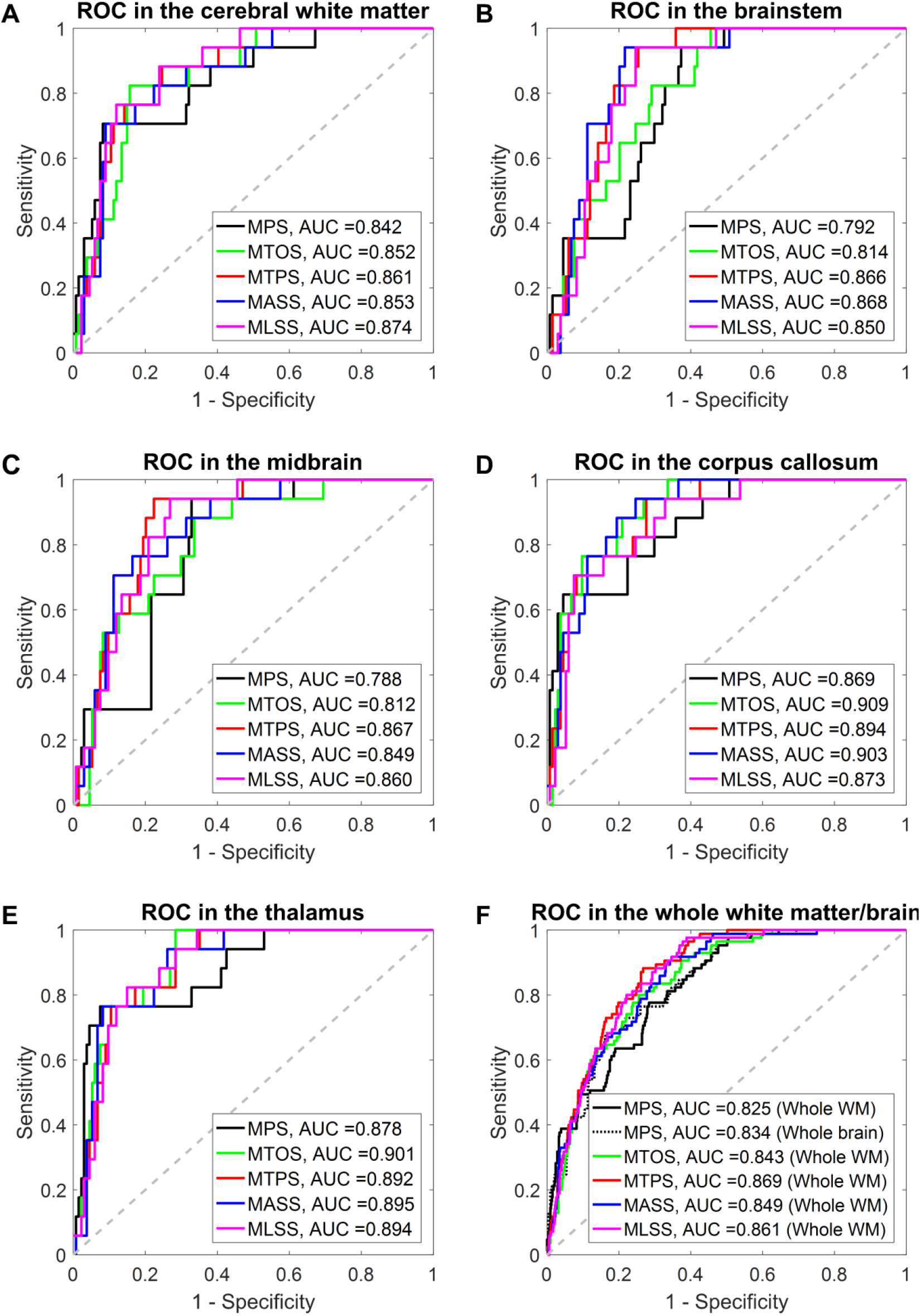
Receiver operator curve (ROC) for the 5 strain-based metrics evaluated at subregional level (**A**-**E**) and the whole white matter/brain level (**F**) based on the testing dataset within the leave-one-out cross-validation framework. The area under the curve is also reported for each metric. In each subfigure, the 50% probability line (gray dash) indicates the level of a random guess (i.e., AUC=0.5).

## Discussions

The current study detailed a theoretical strain analysis to comprehensively quantify the deformation of the WM fiber tracts through coordinate transformations, in which the strain tensor of each WM element was transformed to the coordinate systems with one axis aligned with the real-time fiber orientation. Exact analytical formulas of three novel tract-related strains were presented, characterizing the normal deformation perpendicular to the fiber tracts, and shear deformation along and perpendicular to the fiber tracts, respectively. The injury predictability of these three newly-proposed strain peaks was evaluated by correlating these metrics with injury outcome and compared to that of the previously-used MTOS and MPS. It was verified that four tract-related strain peaks exhibited superior performance in discriminating the concussion and non-concussion cases than the MPS. This comprehensive delineation of the deformation pertaining to the WM fiber tracts suggests the preferential use of tract-related strains as predictors for brain injury.

When focusing on MPS and MTOS, enhanced performance on injury discrimination was noted for MTOS with respect to MPS, correlating well with the findings in previous computational studies.^30, 36, 43, 45, 46, 55^ However, substantial differences existed among the previous studies, particularly on the brain material modeling (e.g., isotropic vs. anisotropic, heterogeneous vs. homogeneous) and the computation of tract-oriented strain (e.g., time-variant fiber orientation vs. time-invariant fiber orientation).^54^ The current study simulated the brain as an isotropic and homogenous structure and leveraged an embedded element approach to monitor the real-time fiber orientation during head impacts. This orientation information was further used to inform the calculation of tract-oriented strain by projecting the strain tensors from pre-computed simulation along the real-time fiber direction. Thus, the current study can be regarded as a significant extension to previous efforts,^30, 36, 43, 45, 46, 55^ collectively further reinforcing the importance of adopting orientation-dependent metrics as predictors for brain injury, especially when it comes to concussion or axonal injury.

Besides the MPS and MTOS, the current study proposed three new tract-related strain peaks as injury metrics, i.e., MTVS, MASS, and MLSS, contributing to a comprehensive delineation of WM tract-related deformation with respect to the previous practice where only the MTOS was presented.^9, 30, 34–37, 39–54, 81, 82^ By correlating the injury metrics with the injury outcome, the three newly-proposed metrics exhibited comparable discrimination performance to the MTOS, but were consistently superior to the MPS. These results were extensively supported by the findings in *in vivo, ex vivo*, and *in vitro* experimental models.^56–59, 83–93^ For example, LaPlaca and colleagues^58^ found that a significant loss of neurites in the *in vitro* model occurred in the regime at which the axial-shear strain peaked. More recently, another *in vitro* model ^56^ also found that the spatial distribution of high axial-shear strains correlated with the localization of neurite blebbing. By imposing uniaxial stretching to a 2D culture slices with the orientation of the embedded axons quantified, Nakadate and coworkers^59^ found swellings were concomitantly noted in these axons with their orientations parallel and perpendicular to the stretching direction, indicating that both the tract-oriented strain and tract-perpendicular strain might instigate axonal injury. Collectively, the congruency between the current study and experimental models further highlights the potential of the three newly-proposed tract-related strains and the previously-used tract-oriented strain as injury predictors compared to MPS. Nevertheless, it is worth clarifying that the scale of interest in the *in vivo, ex vivo*, and *in vitro* models (e.g., axon/neuron/nerve fiber) may not necessarily be the same as the one in our computational results (i.e., fiber tracts).^46^

Among the four tract-related strain metrics, a regional dependency was noted for the best injury predictor. In all regions, the AUC value of the best predictor was only marginally larger than the three other tract-related strain metrics and MPS. Such results might be partially associated with the following three aspects: Firstly, injury severity of the simulated impacts was diagnosed with binary classification (i.e., concussion or non-concussion), lacking definitive documentation of localization and extent of injury pathology. This was further confounded by the fact that no definitive consensus has been reached regarding the clinical criteria for concussion diagnosis.^94^ Recent studies have also supported that concussion does not inform the pathological cascade secondary to exterior insults and cannot be regarded as a true diagnosis.^95^ Thus, the oversimplified injury outcome, as well as associated compounding factors in the concussion diagnosis of the two simulated datasets,^69, 70^ prohibits further discrimination especially among these four tract-related strain metrics. Secondly, the rather low ratio of concussed cases to all simulated cases (11%) posed another hamper for effectively evaluating the predictive power of these five metrics even within the LOOCV scheme.^80, 96^ With the emergence of open-access platforms for measuring and sharing high-quality head impact kinematics across various institutions,^97^ further verification of the findings in the study is hoped to be granted soon based on additional injury dataset with the concussive and non-concussive impacts optimally distributed. Thirdly, the discrimination performance of all metrics was evaluated based on the 95^th^ percentile peak strain values, the same as the strategy in previous studies.^28, 43, 52, 77^ This strategy is a typical down-sampling operation, in which the spatiotemporally-detailed strain information estimated by the FE model was represented by one single variable characterizing the peaking response over time across a pre-defined region. The reliability of such a down-sampling operation and corresponding effects on injury prediction performance remains to be further investigated.^52, 53^ Moreover, as exemplified by one representative element in Fig. 4, the fiber tracts experienced alternate combinations of normal and shear deformation with the predominant mode varied across the timespan of the impact. It might be possible that brain injury was the synergistic outcome of various strains with their effects either alternately or simultaneously exerted. Thus, another possible direction for future work could be identifying the synergism of combined strain measures and corresponding influence on injury discrimination, instead of exclusively relying on only one metric.

Axonal injury is a complex injury with a wide spectrum of pathological severity, ranging from mild levels, which can be associated with subtle neurological deficits pertaining to electrophysiological impairment, to several forms, which can lead to a large scale of neuronal degeneration/death throughout the WM.^98, 99^ Through careful spatiotemporal correlation between specific manifestation of neural pathology and various strain measures, an *in vitro* model recently found that the bleb formation was strongly correlated with axial-shear strain, while being uncorrelated with the tract-oriented strain.^56^ Inversely, the timeframe of neuronal death was only correlated with the integral of tract-oriented strain over the length of the neuron, rather than its counterparts based on the axial-shear strain.^56^ Such results highlighted that the four tract-related strains might have the potential of discriminating different pathological demonstrations. Thus far, the authors are not aware of such an injury dataset with definitive documentation of the location and specific manifestation of injury pathology as well as accurate measurement of the impact loading in humans. A further evaluation of the tract-related strains on their discrimination capability of specific pathological manifestation warrants further work.

As reflected by our modeling choice that no relative motion between the truss elements and their master elements were permitted,^54^ these analytical solutions in Table 1 were attained based on the assumption that the displacements across the fiber tracts and surrounding matrix was continuous, similar to the assumption in the aforementioned *in vitro* models^56–58^ and other computational studies.^34, 35, 45, 46, 51, 82, 100, 101^ This assumption was partially verified by one *in vitro* model, in which no evident movement of soma/neurite with respect to the surrounding matrix was observed in the rat.^58^ However, the authors acknowledge that a systematic examination of the coupling response between the WM fiber tracts and surrounding matrix in humans is deemed necessary.

The presented theoretical framework can also be extendedly implemented to other TBI scenarios, such as mapping strains onto the branched microstructures of the vasculature to uncover the underlying mechanical cue for the blood-brain barrier,^102^ and even far beyond the central nervous system. An extensive discussion regarding the potential implementation of the current theoretical framework can be found in a previous study by Scimone and colleagues.^103^ Another important implication of the current work is to guide the multiscale modeling of axonal injury, which explores the mechanobiological pathway of axonal injury from the macroscopic loads endured at the whole head level to the cellular/molecular level. Current multiscale endeavors often employ a single loading mode, such as uniaxial stretching (corresponding to the tract-oriented strain in the current study).^104–109^ Our work may better inform the multiscale modeling with diverse and combined loadings.

## Other limitations and path forward

Several extra limitations, other than the few highlighted in the “Discussion” section, should be acknowledged which requires further investigation. To ensure computational efficiency both in solving the impact simulations and extracting the tract-related strains, the KTH head model with a mean element size around 5.2 mm was used, in which the embedded fiber orientation was obtained by downsampling the diffusion information from the DTI with a resolution of 1 mm via a weighted average approach. Such an averaging approach inevitably lost orientation information and further affected the computed tract-related strains, since the averaged fiber orientation may not be aligned with the true orientation of the axonal fiber bundles. Consequently, the current endeavor of evaluating the predictive power of four tract-related strains presented in this study should be regarded as a first step and requires further assessment. Compared to the current study where only one fiber orientation was assigned to each WM element, several pioneering studies have more appropriately inserted the fiber orientation from the DTI to the FE models, in which the loss of orientation information was partially alleviated or entirely circumvented. For example, one promising approach to leverage the fiber orientation information from DTI was to explicitly embed the whole WM fiber tractography into the FE head model,^34, 35, 46^ although this approach confronts multiple challenges of tractography reconstruction.^110^ Ji and coworkers^48^ instead transformed the WM voxels and their fiber orientation from the DTI space into the coordinate system of the FE head model. They then computed the tract-oriented strain at the voxel level by resolving the strain tensor of the WM element from pre-computed simulation to the fiber orientation of these voxels that were enclosed by the given WM element. Taken inspiration from the study conducted by Ji and coworkers^48^ in which no averaging was needed, we alternatively suggest that, instead of embedding only one truss element per WM element as is done in the current study, orientation information of all voxels can be wholly integrated into the FE model by embedding multiple truss elements per WM element. Future work can exploit these advanced coupling approaches outlined above to better inform the computation of tract-related strains with their injury predictability being further assessed, especially these three tract-related strains newly proposed in the current study.

Additionally, the orientation information of WM fiber tracts embedded within the FE head model is based on the primary eigenvector of the diffusion tensor. It is well-known that the tensor model is a limited approximation of the complexity of human white matter,^111^ especially to these regions, where many fiber bundles intertwine and cross constituting a significant portion of WM. With the emergence of more sophisticated models^111, 112^ to describe the diffusion in voxels with multidirectional fibers, the current work can be potentially extended by incorporating a full spectrum of orientation information via embedding multiple truss elements per WM element towards an anatomically more realistic description of WM architecture.

Another limitation of the current study is the classification of WM elements. Brain elements with averaged FA value (Equation 2) over 0.2 were considered as WM and further involved in the computation of tract-related strains. This classification protocol distinguished most white matter (e.g., cerebral white matter) from the cortical region (i.e., cortical gray matter). However, it did not exclude the deep brain regions with both gray matter (major component) and white matter (minor component), such as thalamus and midbrain which of both classified as WM in the current study. Fundamentally, this is associated with our strategy of adopting an averaged description of FA value per element (Equation 2), which caused a systematic loss of information relating to the FA. This limitation can be addressed by either exploiting a more advanced coupling approach as described above or adopting a model with a more refined mesh at the deep brain regions.

The current study simulated the brain as a homogenous and isotropic structure, similar to other computational studies.^9, 30, 36, 37, 48^ This is a compromised strategy since no consensus has been reached thus far about the mechanical heterogeneity and anisotropy of the human brain tissue.^63, 113^ For example, Budday and coworkers^14^ reported that the WM was significantly more compliant than the cortical GM with the tissue samples tested under multiple loading modes, while two independent studies^114, 115^ reported the opposite trend using indentation. Controversial findings are also noted about brain anisotropy, particularly for WM. For example, Jin and colleagues^116^ found that mechanical stiffness of WM in shear exhibited directional dependency with respect to the fiber orientation, while mechanical testing performed by two independent groups reported WM was not significantly anisotropic.^14, 117^ Addressing these mechanical ambiguities described above is out of the scope of the current study. Nevertheless, the authors recognize the large number of computational models^34, 35, 39–41, 43, 46, 53, 118^ having incorporated an anisotropic and/or heterogeneous representation of brain material with the exact incorporating approaches being model-specific or application-specific. Therefore, evaluating the predictability of tract-related strains within the context of simulating the brain as an anisotropic and heterogeneous medium is another interesting aspect of future work.

In contrast to previous studies with subject-specific head models being used,^48, 119^ the current study adopted instead a generic head model with the geometry approximately representing the head of 50^th^ percentile male and the fiber orientation extracted from the ICBM atlas to study the cohort of football players. This is due to the unavailability of neuroimaging data of these players, prohibiting a possible integration of subject-specific variability in the current analysis. We recognize that several independent research groups^9, 120–122^ have established technical pipelines facilitating the development of subject-specific FE model, in all of which the displacement field attained via imaging registration was leveraged to inform the model personalization. These personalization pipelines can be integrated with the current study to investigate the deformation of axonal fiber tracts on an individual-specific basis.

Finally, although the current study aims to advance the description of WM tract-related deformation by proposing three new tract-related strain-based metrics, we acknowledge that a complete delineation of WM fiber deformation is yet to be attained. For example, it is expected that, during the head impacts, the fiber tracts may also endure extra deformation modes such as twisting and bending, which remain to be appropriately extracted in the future.

## Conclusion

The present study performed a theoretical strain derivation to comprehensively quantify the deformation pertaining to the WM fiber tracts by relating the strain tensor of the WM element to its embedded fiber tract. In addition to the previously used tract-oriented strain quantifying the normal deformation along the fiber tracts, three new tract-related strains were proposed with exact analytical formulas, characterizing the normal deformation perpendicular to the fiber tracts and shear deformation along and perpendicular to the fiber tracts. By relating the continuous injury criteria and binary injury outcome, it was preliminarily revealed that all tract-related strain peaks demonstrated enhanced discriminating capability of separating the concussion and non-concussion cases than the MPS, advocating the further investigation of tract-related metrics as injury predictors. This study presents a comprehensive delineation of WM tract-related deformation and supports the use of orientation-dependent criteria with injury criteria, which may ultimately contribute to an advanced mechanobiological understanding and enhanced computational predictability of axonal injury.

## Conflict of Interest

Drs. Michael Zeineh, Gerald Grant, and David Camarillo received funding from the Pac-12 Conference’s Student-Athlete Health and Well-Being Initiative and Taube Stanford Children’s Concussion Initiative. Drs. Svein Kleiven and Xiaogai Li received funding from the Swedish Research Council (VR-2016-05314 and VR-2016-04203), while Dr. Marios Georgiadis received funding from the Swiss National Science Foundation (P400PM_180773). The content of this article is solely the responsibility of the authors and does not necessarily represent the official views of funding agencies. The simulations were performed on resources provided by the Swedish National Infrastructure for Computing (SNIC) at the center for High Performance Computing (PDC). The authors also thank Dr. Annaclaudia Montanino for the valuable sharing of literature knowledge and the anonymous reviewers for the stimulating comments and valuable suggestions that substantially improved this paper.

## Appendix

**Fig. A1.**
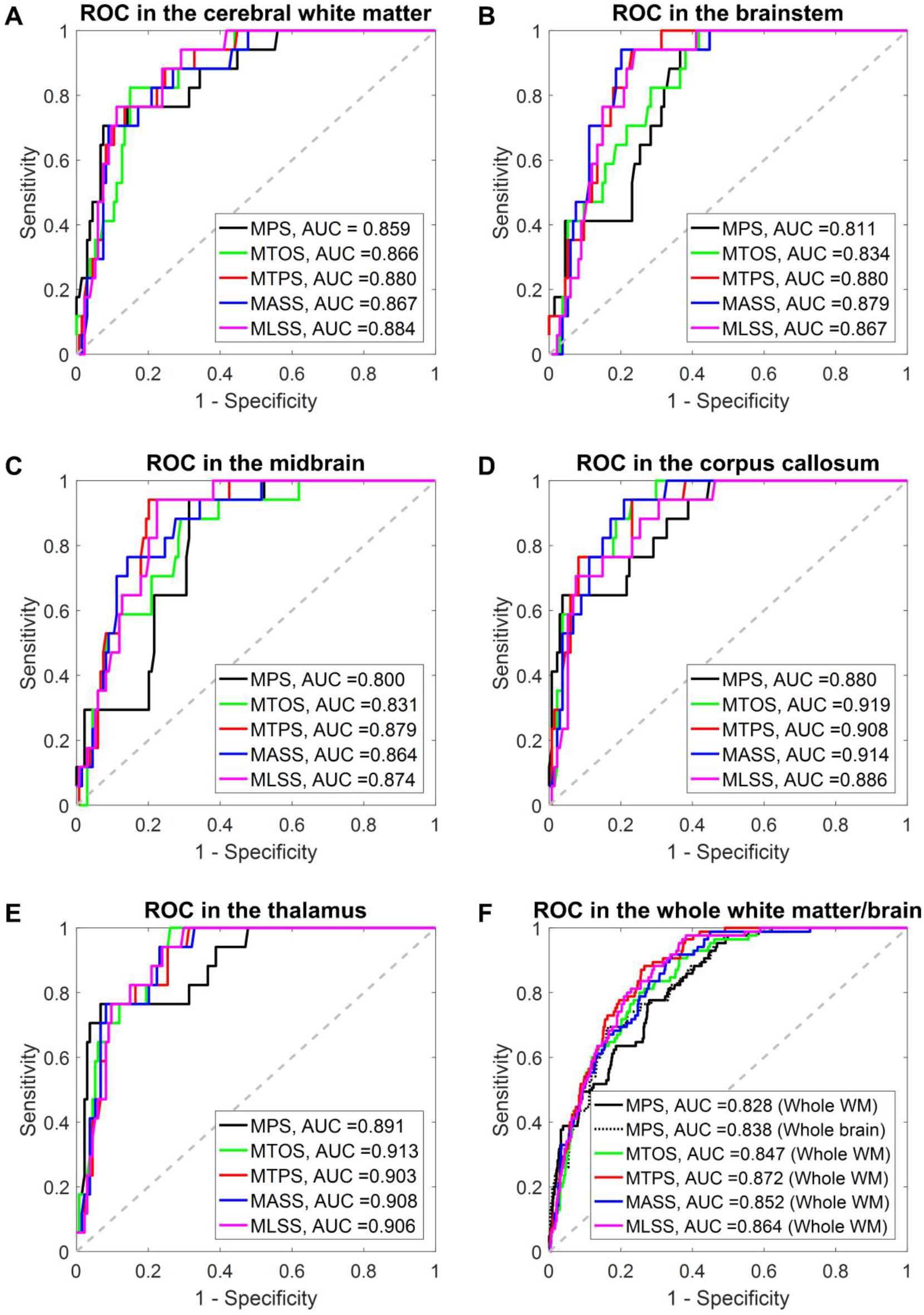
Averaged receiver operator curve (ROC) for the 5 strain-based metrics evaluated at subregional level (**A-E**) and the whole white matter/brain level (**F**) based on the training datasets within the leave-one-out cross-validation framework. The area under the curve is also reported for each metrics. In each subfigure, the 50% probability line in gray dash is additionally plotted, indicating the level of a random guess (i.e., AUC=0.5).

## Notes

### Competing Interest Statement

The authors have declared no competing interest.

